# High-throughput screening of the ReFRAME library identifies potential drug repurposing candidates for *Trypanosoma cruzi*

**DOI:** 10.1101/2019.12.11.873711

**Authors:** Jean A. Bernatchez, Emily Chen, Mitchell V. Hull, Case W. McNamara, James H. McKerrow, Jair L. Siqueira-Neto

**Affiliations:** Skaggs School of Pharmacy and Pharmaceutical Sciences, University of California, San Diego, La Jolla, California, USA; Center for Discovery and Innovation in Parasitic Diseases, University of California, San Diego, La Jolla, California, USA; Calibr, a division of The Scripps Research Institute, La Jolla, California, USA

**Keywords:** *Trypanosoma cruzi*, antiparasitics, high-throughput screening, drug repurposing

## Abstract

Chagas disease, caused by the kinetoplastid parasite *Trypanosoma cruzi*, affects between 6 and 7 million people worldwide, with an estimated 300,000 to 1 million of these cases in the United States. In the chronic phase of infection, *T. cruzi* can cause severe gastrointestinal and cardiac disease, which can be fatal. Currently, only benznidazole is clinically-approved by the FDA for pediatric use to treat this infection in the USA. Toxicity associated with this compound has driven the search for new anti-Chagas agents. Drug repurposing is a particularly attractive strategy for neglected diseases, as pharmacological parameters and toxicity are already known for these compounds, reducing costs and saving time in the drug development pipeline. Here, we screened ~ 12,000 compounds from the ReFRAME library, a collection of drugs or compounds with confirmed clinical safety, against *T. cruzi*. We identified 7 compounds of interest with potent *in vitro* activity against the parasite with a therapeutic index of 10 or greater, including the previously-unreported activity of the antiherpetic compound 348U87. These results provide the framework for further development of new *T. cruzi* leads that can potentially move quickly to the clinic.

## Introduction

*Trypanosoma cruzi*, the causative agent of Chagas disease, is a protozoan parasite that is primarily transmitted to humans via triatomine insects (known as kissing bugs) during a blood meal. Infection by *T. cruzi* manifests initially in an acute phase of infection, and if left untreated, proceeds to a chronic phase (1). During the acute phase, mild or unremarkable symptoms such as fever, fatigue, rash, headache or swelling at the site of the triatome bite may present. When left untreated, the primary infection usually resolves in weeks, but residual parasites remain in the host’s body evolving to the chronic phase. Over the span of years to decades, approximately 30% of those infected individuals will manifest cardiac and/or gastrointestinal complications leading to morbidity and mortality (2).

Current treatment options are very limited for Chagas disease: only benznidazole is clinically-approved for pediatric use in the case of acute *T. cruzi* infections in the United States. Benznidazole and nifurtimox are available off-label via the CDC for compassionate use for all other cases of this infection. However, severe side-effects associated with the use of these medications leads to high levels of patient discontinuation of treatment. Furthermore, the usefulness of benznidazole in the chronic phase of infection is disputed within the Chagas research community (3–5).

Limited efforts from the pharmaceutical industry to develop a medication for *T. cruzi* infections further complicates progress towards anti-Chagas agents better than benznidazole and nifurtimox. Rising costs and high levels of failure of drug molecules in clinical trials due to adverse events and lack of efficacy present further general barriers to the development of medications. One cost-effective strategy involves repurposing existing drugs with known toxicity and pharmacokinetic profiles for other indications (6). This has the potential to speed up drug development efforts, reduce costs and lower the chance of adverse events presenting in clinical trials. The ReFRAME (<u>Re</u>purposing, <u>F</u>ocused <u>R</u>escue, and <u>A</u>ccelerated <u>Me</u>dchem) library, a comprehensive set of molecules with tested clinical safety, has been previously used to identify potential drug repurposing hits for neglected tropical diseases (7, 8). In this work, we screened 7,680 compounds from this library against the medically relevant intracellular, amastigote form of *T. cruzi* infecting mouse myoblasts using a high-throughput, phenotypic cellular imaging assay that our group has successfully used in previous studies to identify novel antitrypanosomal agents (9–11). We identified seven compounds with suitable selectivity indexes (SIs) for drug repurposing; two of these, the antiherpetic drug 348U87 and the serotonin receptor binder 3-[4-[4-(2-Methoxyphenyl)piperazine-1-yl]butyl]-6-[2-[4-(4-fluorobenzoyl)piperidine-1-yl]ethyl]benzothiazole-2(3H)-one, have not previously been reported as anti-Chagas compounds and may target the parasite through a novel mechanism. These molecules form an attractive collection of lead molecules for potential drug repurposing to treat Chagas disease.

## Results

### Primary screening of the ReFRAME library against T. cruzi using a high-content imaging assay

Compounds from the ReFRAME library were pre-spotted on 1536 clear-bottom black well plates in 100% dimethylsulfoxide (DMSO) for a final concentration of 10 µM in 10 µL final volume (and 0.1% DMSO final concentration). C212 mouse myoblasts and CA-I/72 strain *T. cruzi* trypomastigotes were added to the plate in a 15:1 infection ratio and incubated for 72 hours at 37°C and 5% CO_2_. Cells were then fixed with 4% paraformaldehyde (final concentration), and stained with 5 µg/ml of 4’,6-diamidino-2-phenylindole (DAPI) to highlight the nuclei from the host cells and parasites. Using an ImageXpress MicroXL automated microscope (10x magnification setting), fluorescence images of C2C12 and *T. cruzi* amastigote nuclei were acquired for each well, and the number of host cell nuclei and amastigote nuclei were automatically determined using a custom image analysis module as previously described (9–11) (Figure 1). Infection levels were calculated as a ratio of the number of *T. cruzi* amastigotes per C2C12 host cell as determined by nuclei counting. Compound toxicity was determined by dividing the number of host cell nuclei in a drug-treated well to the average of the vehicle controls. For both infection ratios and cell viability ratios, values were normalized to the vehicle controls to determined percent activity and toxicity, respectively. Control wells containing uninfected C2C12 cells, infected C2C12 cells with 0.1% DMSO, and infected C2C12 cells with 0.1% DMSO and 50 µM benznidazole were prepared in each plate for data normalization. The mean Z’ for the 10 plates tested was 0.52 with a standard error of 0.01. Hit selection cutoffs for the primary screen were set at 70% antiparasitic activity (70% parasite reduction compared to untreated controls) and 50% host cell viability at 10 µM compound compared to untreated controls (represents approximately 3 standard deviations from the average of the untreated controls). We identified 238 compounds (2% of the total library) that met these selection criteria. A summary of the primary screening data is shown in Figure 2.

**Figure 1.**
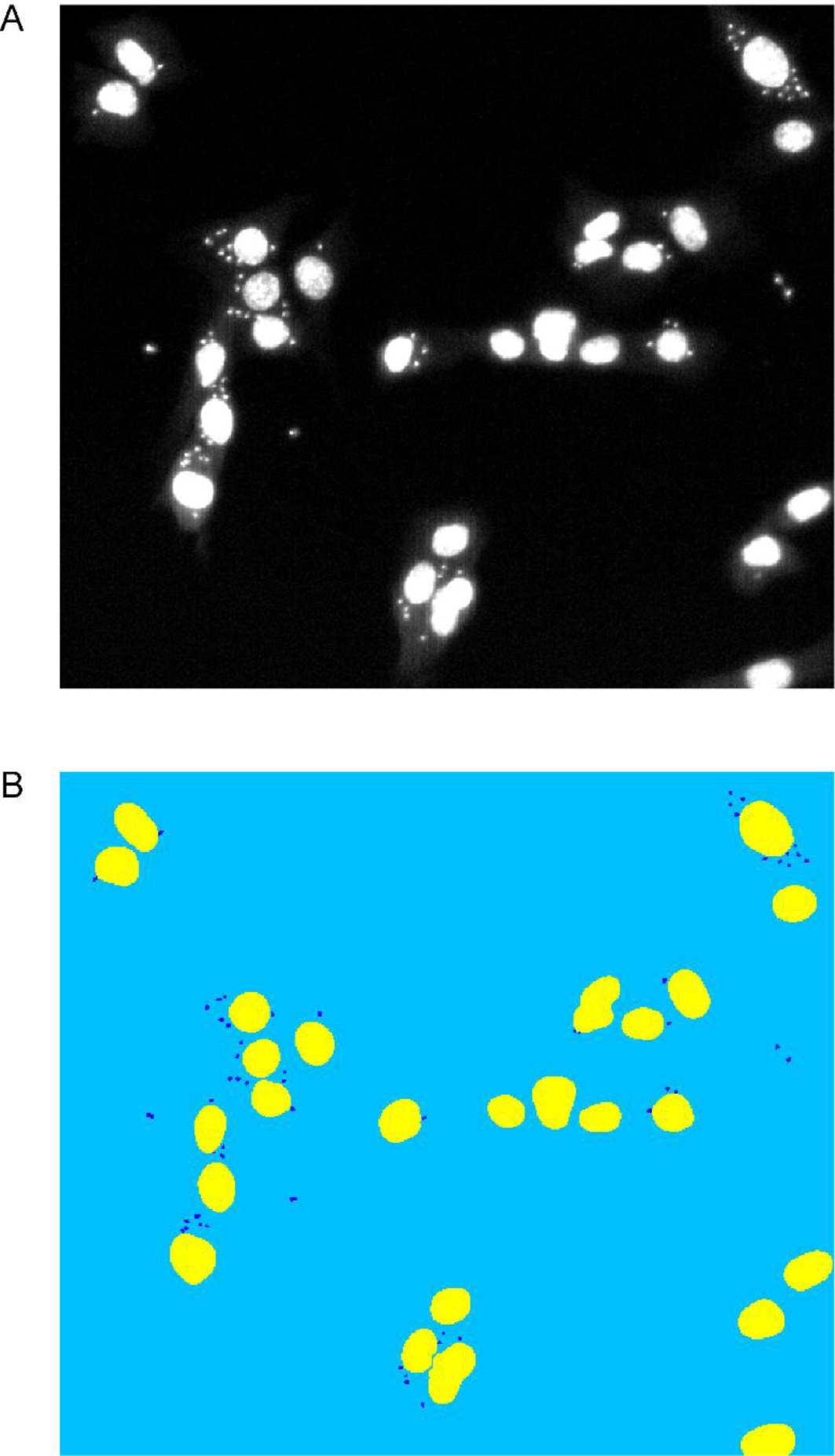
Automated segmentation analysis of *T. cruzi* amastigote and C2C12 mouse cardiomyocyte nuclei. A) Fluorescence microscopy image (10×magnification) of DAPI-stained C2C12 infected with CA-I/72 *T. cruzi* amastigotes 72 hours post-infection. B) Custom module segmentation of host and parasite cell nuclei using MetaXpress 5.0 (Molecular Devices). Host cell nuclei are in yellow and parasite nuclei are in dark blue.

**Figure 2.**
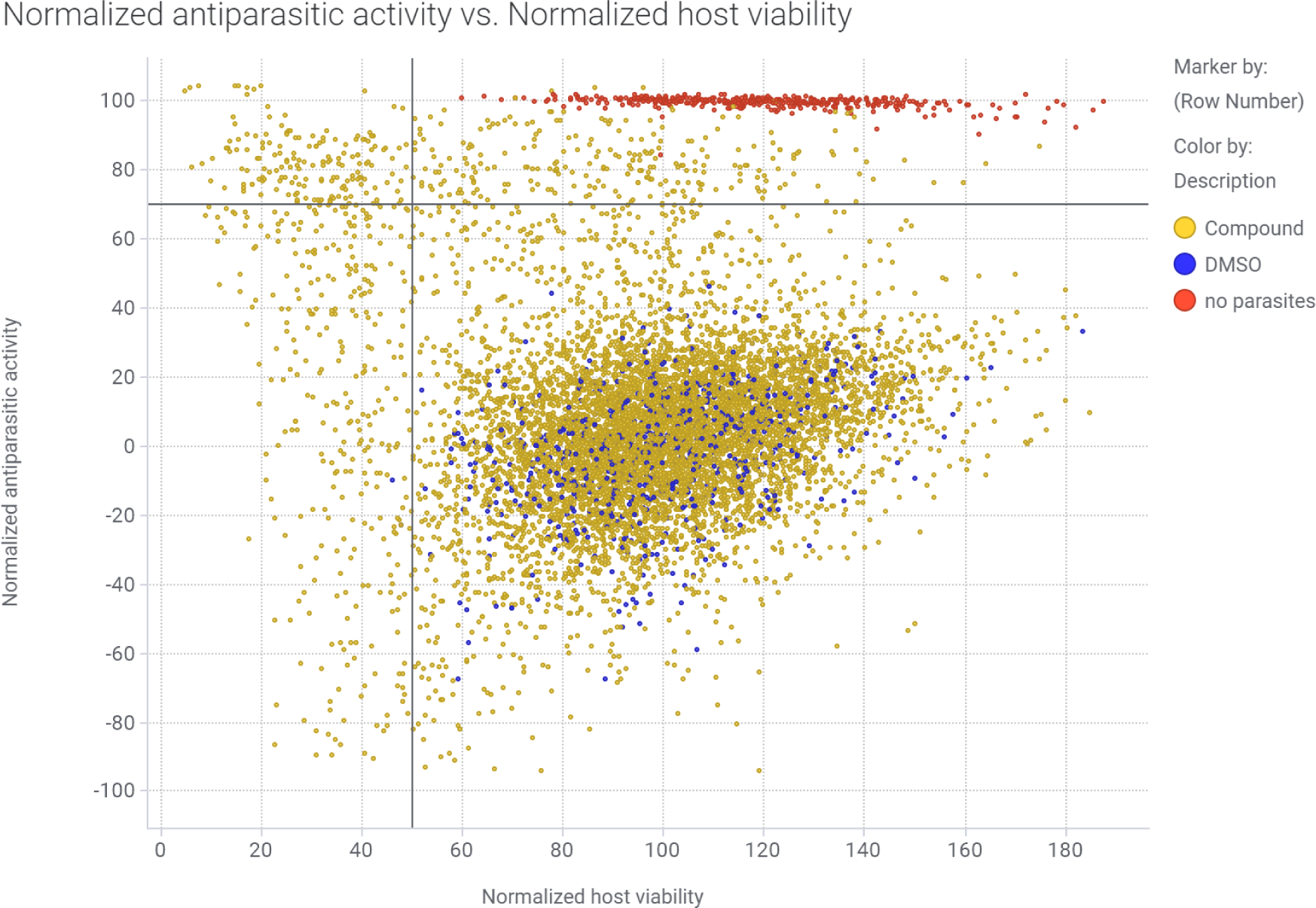
Primary screening data for the ReFRAME library against *T. cruzi* in the phenotypic high-content imaging assay. Scatter Plot of normalized % activity against CA-I/72 *T. cruzi* (Normalized antiparasitic activity %, Y-axis) and the host cell C2C12 viability % (Normalized host viability %, X-axis) for ReFRAME library. The red dots represent the uninfected controls, blue dots represent untreated controls (0.1% DMSO) and the yellow dots are the tested compounds. A vertical line at 50% Normalized host viability and a horizontal line at 70% Normalized antiparasitic activity highlights the top right quadrant where the compounds were selected as hits for dose-response confirmation.

### Counter-screen of 238 hits in dose response

To validate the hits obtained from our primary screen, compounds were re-spotted in duplicate in a 10-point, 3-fold dilution dose response, with 10 µM as the highest concentration of inhibitor. Using the high-content imaging assay, 238 primary hits were retested in duplicate. We identified seven compounds of interest using the following cutoff criteria: at least 70% antiparasitic activity at a given drug concentration in one of the two dose response replicates and a therapeutic index of 10 or greater. Half maximal effective concentration (EC_50_), half maximal cytotoxic concentration (CC_50_) and selective index (SI; defined by the ratio of CC_50_ to EC_50_) for validated hits are shown in Table 1, their dose response curves are shown in Figure 3 and the chemical structures of these compounds are displayed in Figure 4. The 7 compounds retained were NSC-706744, 348U87, ASP-8273, XR 5944, Prenyl-IN-1, 3-[4-[4-(2-Methoxyphenyl)piperazine-1-yl]butyl]-6-[2-[4-(4-fluorobenzoyl)piperidine-1-yl]ethyl]benzothiazole-2(3H)-one and incadronate disodium.

**Table 1.**
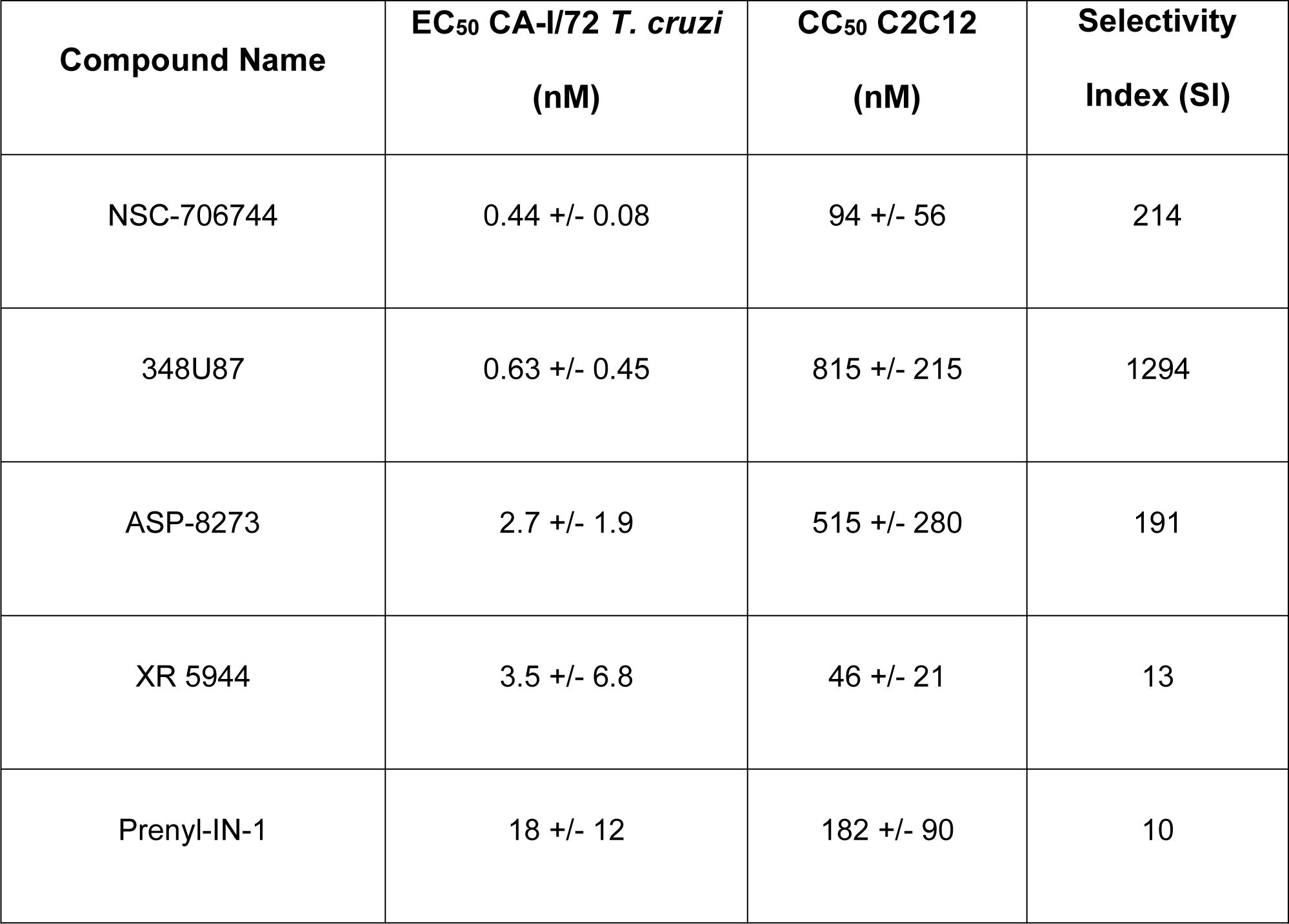
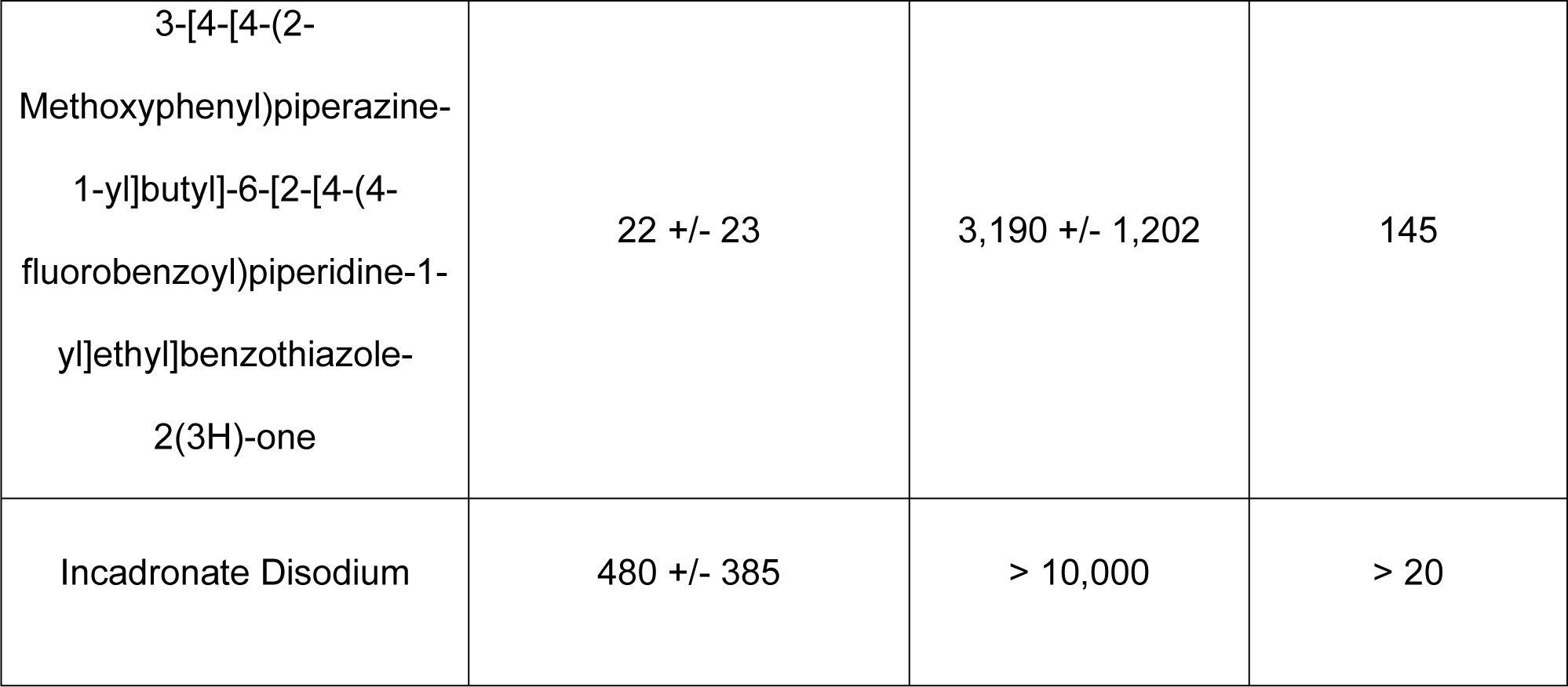
EC_50_ and CC_50_ values for 7 validates hits from the ReFRAME library against CA-I/72 *T. cruzi* in the phenotypic high-content imaging assay ranked in order of potency. Values were calculated from duplicate dose response data, +/− standard error (SE). EC50 values are for *T. cruzi* CA-I/72 parasites. CC_50_ values are for C2C12 cardiomyocyte host cells. Selectivity index = CC_50_/EC_50_.

| **Compound Name** | **EC<sub>50</sub> CA-I/72 *T. cruzi***<br>**(nM)** | **CC<sub>50</sub> C2C12**<br>**(nM)** | **Selectivity**<br>**Index (SI)** |
| --- | --- | --- | --- |
| NSC-706744 | 0.44 +/- 0.08 | 94 +/- 56 | 214 |
| 348U87 | 0.63 +/- 0.45 | 815 +/- 215 | 1294 |
| ASP-8273 | 2.7 +/- 1.9 | 515 +/- 280 | 191 |
| XR 5944 | 3.5 +/- 6.8 | 46 +/- 21 | 13 |
| Prenyl-IN-1 | 18 +/- 12 | 182 +/- 90 | 10 |
| 3-[4-[4-(2-Methoxyphenyl)piperazine-1-yl]butyl]-6-[2-[4-(4-fluorobenzoyl)piperidine-1-yl]ethyl]benzothiazole-2(3H)-one | 22 +/- 23 | 3,190 +/- 1,202 | 145 |
| Incadronate Disodium | 480 +/- 385 | > 10,000 | > 20 |

**Figure 3.**
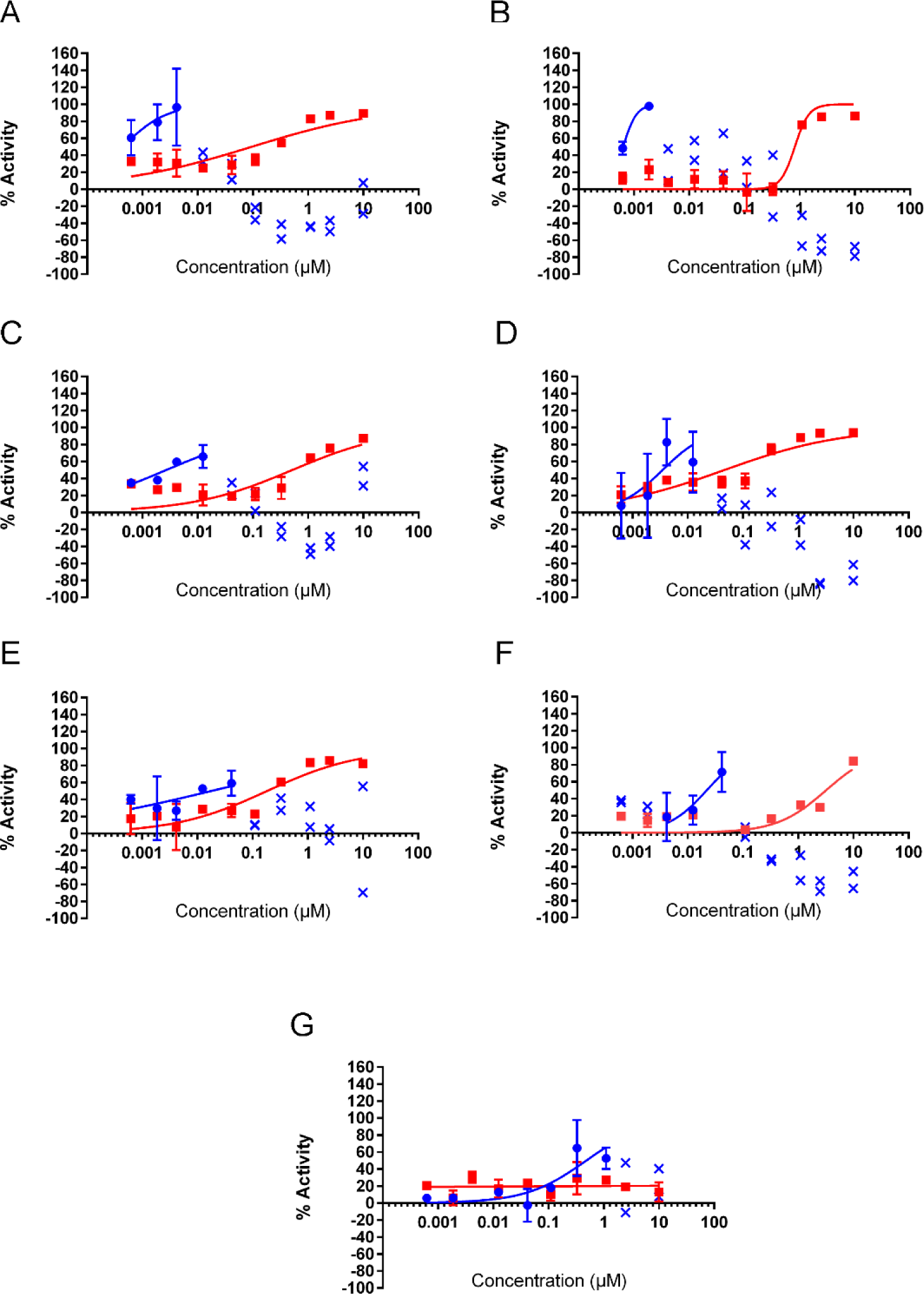
Dose response curves for validated hits from the ReFRAME library. % activity of the hit compounds NSC-706744 (A), 348U87 (B), ASP-8273 (C), XR 5944 (D), Prenyl-IN-1 (E), 3-[4-[4-(2-Methoxyphenyl)piperazine-1-yl]butyl]-6-[2-[4-(4-fluorobenzoyl)piperidine-1-yl]ethyl]benzothiazole-2(3H)-one (F) and Incadronate Disodium (G) against *T. cruzi* (blue curves, antiparasitic activity) and C2C12 cardiomyocytes (red curves, host cell toxicity) are shown for 2 replicate dose responses. Error bars indicate the standard error for each data point. Data values marked in blue with an “x” represent excluded antiparasitic activity data points (where host cell toxicity results in a drop-off of antiparasitic activity).

**Figure 4.**
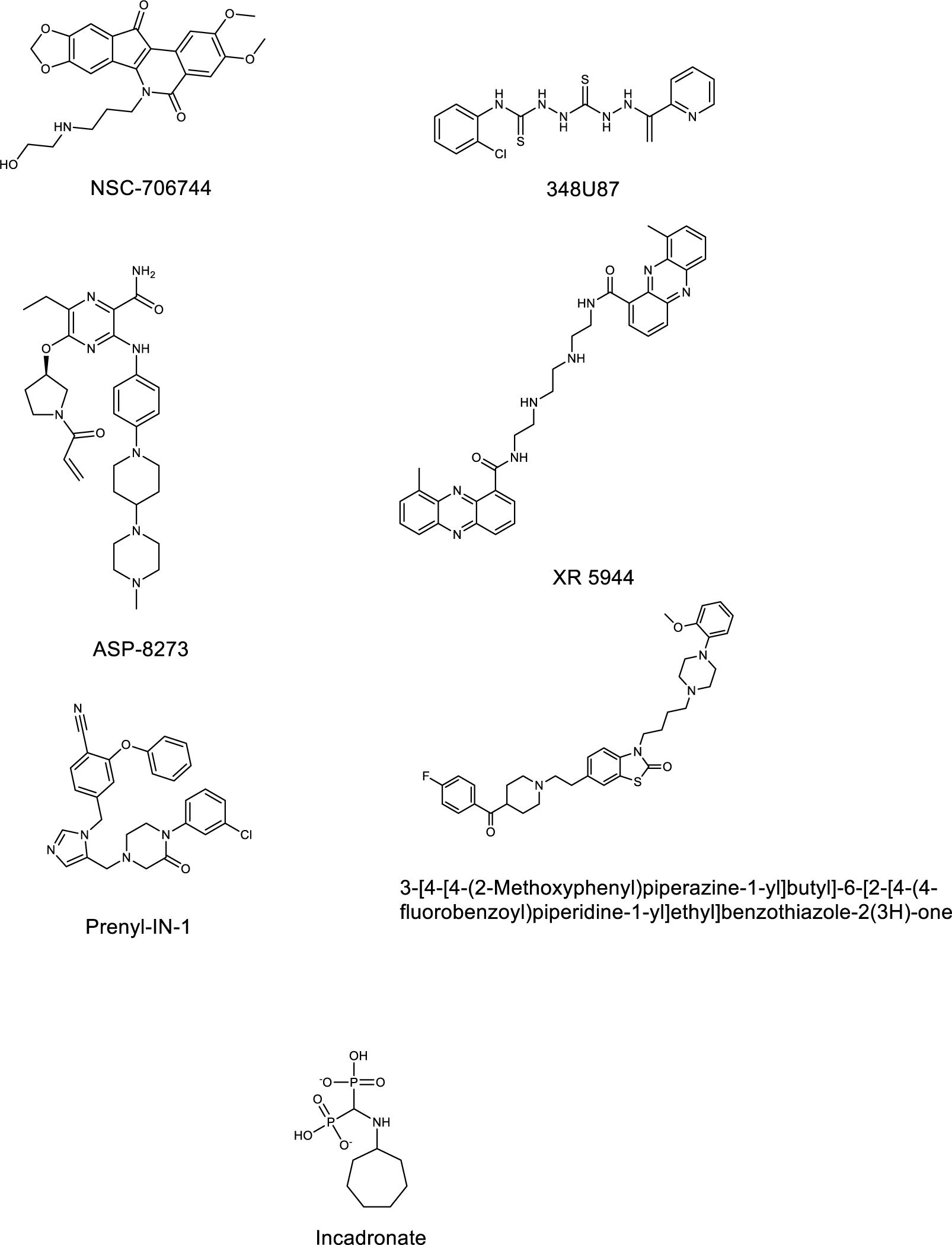
Chemical structures of hit compounds from the high-throughput screening campaign.

## Discussion

Given the toxicity of currently available drugs to treat *T. cruzi* infection, research efforts have been made to explore additional candidate molecules against the parasite. In this work, we screened a drug repurposing library for compounds with anti-Chagasic activity. We identified 7 molecules as having potent *in vitro* activity against *T. cruzi* and a SI of at least 10 against C2C12 cardiomyocyte host cells.

As validation for our assay, we identified a number of compounds from classes which have previously been reported as having anti-Chagas activity, namely farnesyltransferase inhibitors (12–16) (Prenyl-IN-1, incadronate disodium) and DNA topoisomerase inhibitors (17–20) (NSC-706744, XR 5944).

We also identified the EGFR inhibitor ASP-8273 (naquotinib) (21–23) as an inhibitor of *T. cruzi* with good potency (EC_50_ of 2.7 nM) and a SI of 191. Several studies have successfully explored kinase inhibitors of trypanosomatids as therapeutic agents (24–26), and this compound may represent yet another possible candidate for repurposing or further chemical derivatization.

Interestingly, we identified two compounds with intriguing primary indications that may target *T. cruzi* by novel mechanisms. First, serotonin receptor ligands such as 3-[4-[4-(2-Methoxyphenyl)piperazine-1-yl]butyl]-6-[2-[4-(4-fluorobenzoyl)piperidine-1-yl]ethyl]benzothiazole-2(3H)-one (27) have been investigated as agents to treat anxiety and panic disorders. This compound had an EC_50_ of 22 nM and a SI of 145, making it an attractive candidate for drug repurposing and animal model testing. Second, the herpes virus drug 348U87 (EC_50_ of 0.63 nM and a SI of 1294), which targets the viral ribonucleotide reductase (28–31), has been shown to potentiate the activity of acyclovir in topical applications. The thiosemicarbazone iron chelator 3-AP has also been shown to inactivate the ribonucleotide reductase of *T. brucei* (32), and 348U87 may act against a homologous protein of *T. cruzi* in a similar manner. For both of the afore-mentioned compounds, mechanistic studies are planned and ongoing to identify the precise molecular targets of these inhibitors.

Follow-up studies in our group will test our most promising compounds in mouse models of *T. cruzi* infection to establish proper dosing protocols and *in vivo* efficacy. In sum, these compounds represent new potent new leads with known pharmacological parameters and possibly novel mechanisms of action against *T. cruzi*, making them attractive candidates for accelerated development as anti-Chagas agents.

## Materials and Methods

### Cells

C2C12 mouse myoblasts (ATCC CRL-1772) and CA-I/72 *T. cruzi* (kindly donated by J. Dvorak, NIH) were cultured in Dubelco’s Modified Eagle Medium (Invitrogen, 11095-080) supplemented with 5% fetal bovine serum (Sigma Aldrich, F2442) and 1% penicillin-streptomycin (Invitrogen, 15140122) at 37°C and 5% CO_2_ essentially as described (11). Passaging of CA-I/72 *T. cruzi* was conducted weekly via co-culture with C2C12 host cells.

### Phenotypic imaging assay

Compounds from ReFRAME library, benznidazole (Sigma cat. no. 419656) and DMSO (Sigma cat. no. D2650) were transferred to black 1536-well plates (Greiner Bio One, 782092) with clear bottoms using an Acoustic Transfer System (ATS) instrument (EDC Biosystems). C2C12 cells were seeded at a density of 100 cells per well and CA-I/72 *T. cruzi* parasites were seeded at a density of 1,500 cells per well using a Multidrop Combi liquid handler (Thermo Scientific). Plates were incubated at 37°C and 5% CO_2_ for 72 hours in humidified trays to reduce edge effect. Following this incubation, paraformaldehyde (4% final concentration) in 1× phosphate buffered saline (PBS, Invitrogen, 10010023) was used to fix the cells for 1 hour. The cells were then subsequently treated with 5 µg/mL DAPI staining solution (Sigma Aldrich, D9542) for 1 hour. Next, the plates were imaged using an ImageXpress Micro automated high-content imager (Molecular Devices) using the 10× fluorescence objective. Images were analyzed automatically using a custom image analysis module (9, 11).

### Software

Chemical structures were prepared using ChemDraw Professional 18.1 (Perkin Elmer). EC_50_ and CC_50_ curves were generated using GraphPad Prism 8 (GraphPad Software).

## Acknowledgments

This work was supported by the Bill & Melinda Gates Foundation (OPP1107194). The screening experiments were performed at the UCSD Screening Core.

